# Silymarin Inhibits In Vitro SARS-CoV-2 Infection In Vero E6 Cells

**DOI:** 10.1101/2023.04.07.535766

**Authors:** Nnaemeka Emmanuel Nnadi, Pam Dachung Luka, Simeon Omale, Nathan Yakubu Shehu, John Chinyere Aguiyi

**Affiliations:** Department of Microbiology, Faculty of Natural and Applied Sciences, Plateau State University, Bokkos, Nigeria; Biotechnology Centre, National Veterinary Research Institute, Vom. Nigeria; Department of Pharmacology and Toxicology, Faculty of Pharmaceutical Sciences, University of Jos, Jos, Nigeria; Department of Infectious Medicine, Jos University Teaching Hospital Jos, Jos, Nigeria

## Abstract

The study evaluated the invitro ability of Silymarin to inhibit SARS-COV-2 infection on Vero Cells. We set out to evaluate the hypothesis that Silymarin has both preventive and curative against SARS-COV-2. To study this, we first evaluated the safety profile of Silymarin using the Drosophila melanogaster(Harwich strain) model. Silymarin tablet film coated 140mg(Silybon-140) was used for the study. We evaluated the fly for acute toxicity, Locomotor performance, estimation of total thiol level, determination of Acetylcholinesterases (AchE) activity, Catalase activity, Glutathion-S-transferase(GST) activity and fecundity assay. To evaluate the invitro activity of Silymarin against SARS-COV-2, SARS-COV-2 isolates from oropharyngeal swabs and confirmed using qRT-PCR were cultured in Vero E6 monolayer cells. Different concentrations of silymarin concentration were used to determine pre- or post-exposure activity.

The result showed that daily exposure to silymarin dose between 50% to 2000% adult dose showed no adverse effect after 28 days. Treatment of Vero cells with silymarin at the concentration 250-500ug/ml all revealed a pre-treatment effect to SARS-COV-2 in vitro and no inhibition effect was observed when the virus was first added before the addition of Silymarin. Silymarin had no adverse effect on D. melanogaster and can be used a preventive drug against SARS-COV-2

## Introduction

Drug therapy option against SARS-CoV-2 is limited. To date, no specific drug has been identified to inhibit and control SARS-CoV-2 in COVID-19 patients^1,2^. The rapid and widespread spread of SARS-CoV-2 and some patients’ lethal pneumonia makes it imperative to find effective treatments immediately. To find alternative drugs with effects against the virus, computational methods were used in several studies to identify possible drugs with therapeutic effects to be chosen to identify special antiviral drug candidates ^3-5^. Molecular docking is a vital bioinformatics modelling tool for drug discovery that predicts the “best-fit” intermolecular binding between a small molecule and a target or two proteins at the atomic level. It characterizes ligand behaviour in target protein binding sites and elucidates fundamental biochemical processes^6^

Only a few studies have gone beyond computational to invitro analysis of identified potential drug targets^7,8^. Computational drug design is meant to identify potential drugs in which the best compounds should then be tested experimentally in the lab to ensure their true binding and effectiveness (such as stopping viral infectivity)^9^. We ealier demonstrated silybinin as a potential drug with multi-target interaction with both the S glycoprotein and Mpro protein targets of SARS-COV-2^3^. We hypothesized that silybinin contained in silymarin could be used for drug repurposing in the treatment of COVID-19 disease. In this study, we set out to determine the ability of Silymarin to inhibit SARS-COV-2 on Vero cells, as well as the toxicological and overall safety profile of Silymarin using *Drosophila melanogaster model*.

## Methods

### Toxicity profile

#### Maintaining *Drosophila melanogaster* (Harwich strain)

*Drosophila melanogaster* (Harwich strain) was obtained from the Africa Centre of Excellence in Phytomedicine Research and Development (ACEPRD) Drosophila Research Laboratory at the University of Jos in Plateau state, Nigeria. All flies were kept in vials containing standard fly food at a constant temperature of 25 °C and 70% relative humidity with a 12-hour light/dark clock cycle. All experiments were carried out in accordance with the internationally accepted principles for laboratory animal use and care outlined in EEC directives 1986; 86/609/EEC.

#### Experimental Design

Silymarin tablets film-coated 140mg (silybon-140) manufactured by Micro Labs Limited, 92 Spicot, Hosur-635126, India, with the manufacturing date Feb. 2019 and Exp.Jan. 2022and batch number SYFH0105, was used for the study between September 2020 and February 2021. Acute toxicity experiment was performed for 7 days by exposing young flies of 1-3 days old (both gender) to a wide range of 50% to 2,000% human Adult dose (AD) concentration of silymarin. The mortality rate was monitored, the LD_50_ was calculated and expressed as %survival (fig.1). To assess the drug’s chronic toxicity profile, two to three-day-old flies were exposed to a graded concentration of Silymarin for 28 days (fig.2). The correct concentration of Silymarin was determined and used for the entire seven-day toxicological experiment (fig.3&4). After 7 days, the flies were harvested and homogenized for biochemical assays. Adult dose (AD) reference values were 50% (0.01 mg), 100% (0.02mg), 200% (0.04mg), 500% (0.1mg), 1000% (0.2mg), and 2000% (0.4mg) per 10g diet respectively.

**Fig 1:**
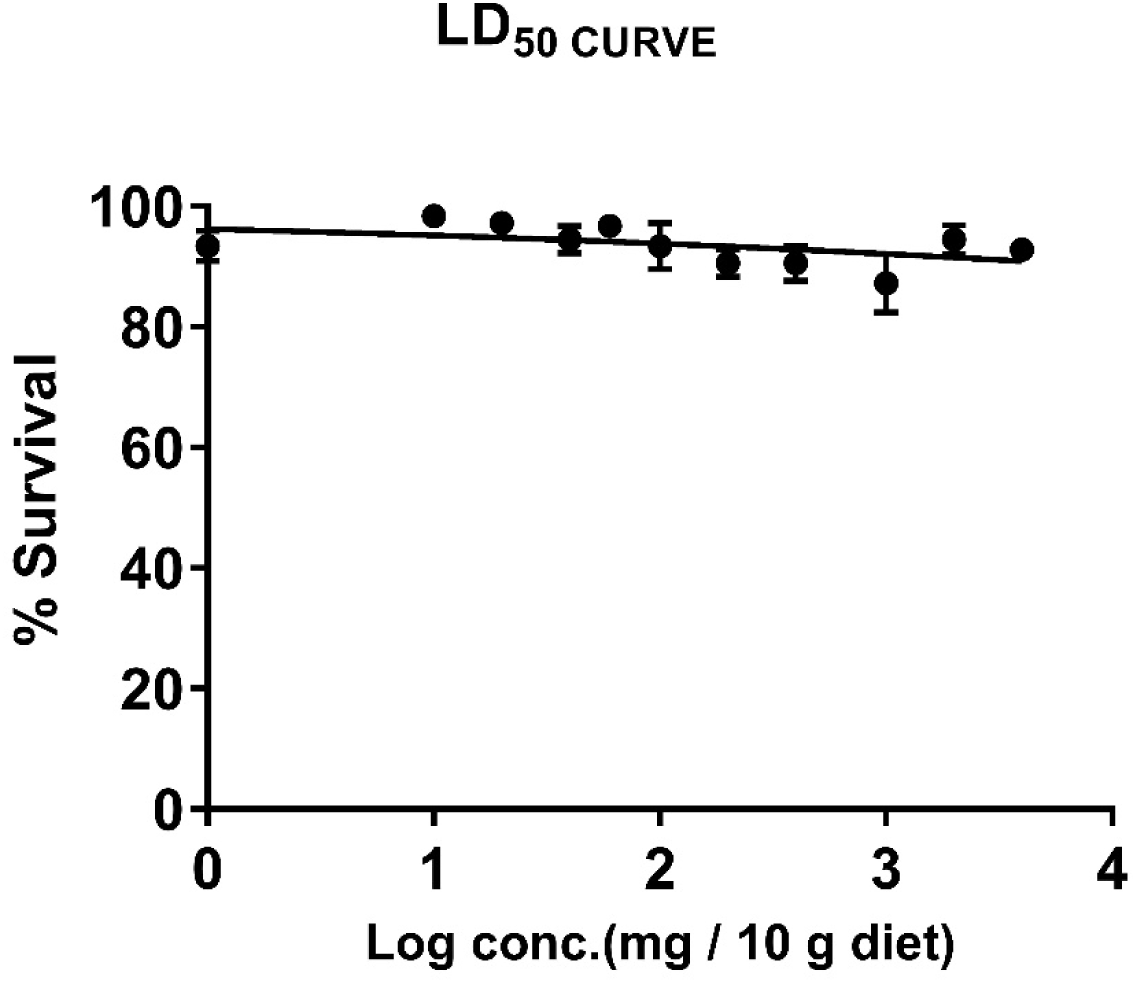
LD_50_ Showing 7-Days Fly Survival at a Concentration Range of 50% to 2,000% Adult Dose (AD) of Silymarin. The results showed that Silymarin is practically safe in the *Drosophila melanogaster* model. Statistical analysis showed mean LD_50_ of above 12,000 mg/10g diet

**Fig 2:**
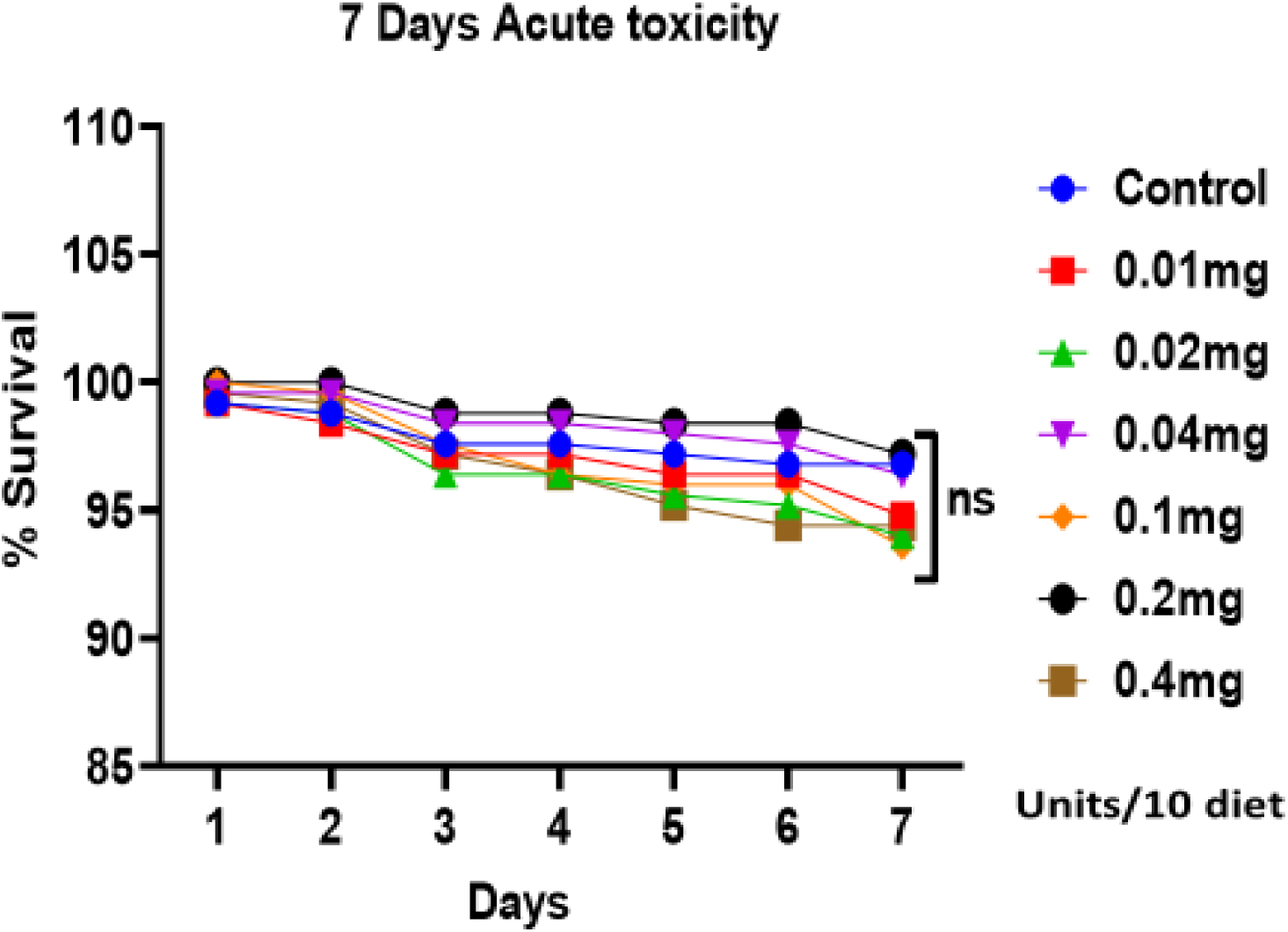
Showing 7- Survival of Daily Oral Dose of Silymarin Tablet in *Drosophila melanogaster* (DM) There was no significant difference (*P> 0*.*05*) between the various groups and the control. A dose of 2000% AD showed no difference in mortality rates compared to the control. *P<0*.*05* *= Statistically significant

**Fig 3:**
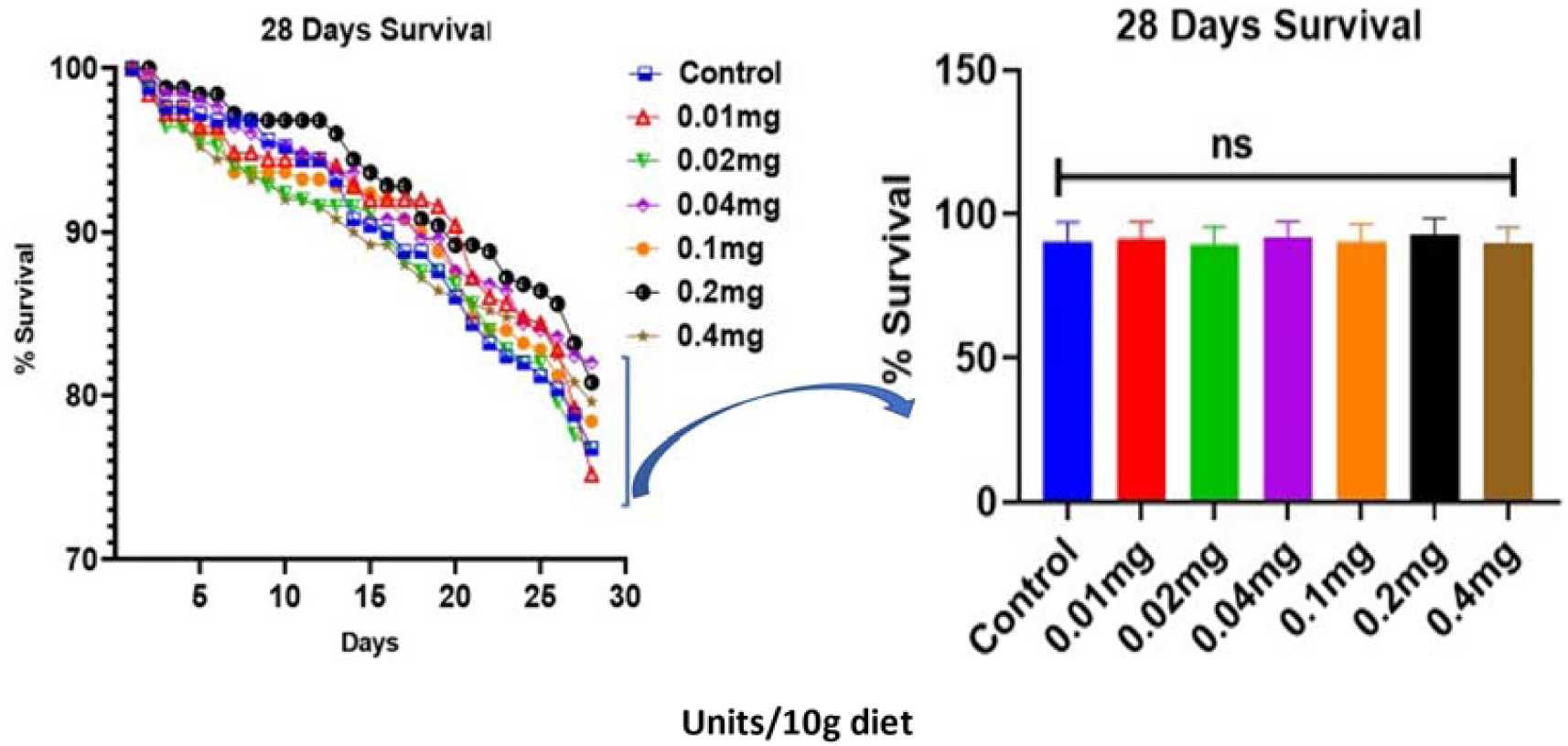
Showing 28-Days Survival Assay in DM. The results showed no significant difference (*P>0*.*001*) between the control and the treated groups. There is also no considerable difference between flies fed with up to 2000 % and 50% oral adult dose of Silymarin. *P<0*.*05* *= Statistically significant

**Fig 4:**
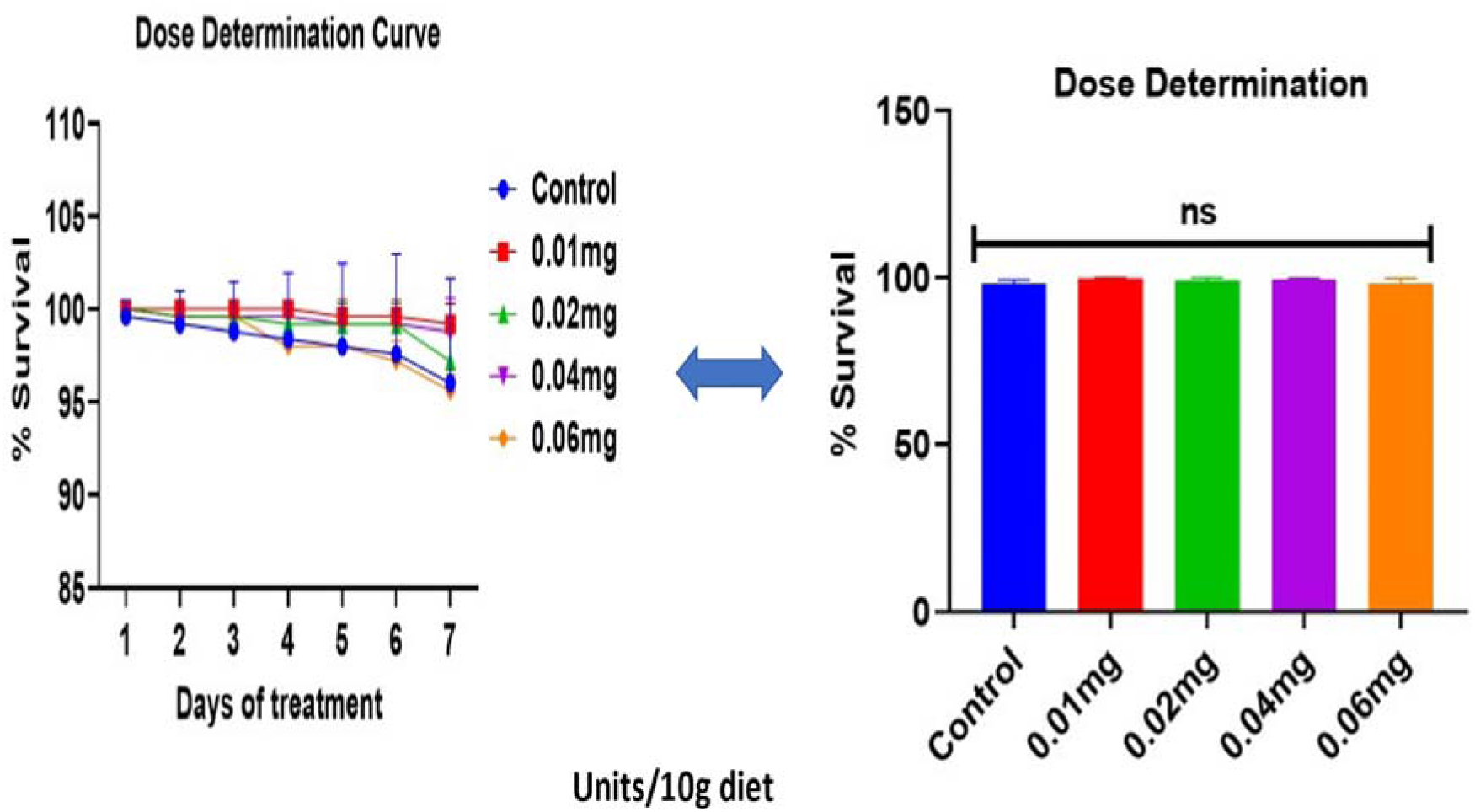
Showing Treatment Dose Determination Curve. The result showed practical safety and there was no significant difference (*P>0*.*05*) for the dose employed in the experiment. The doses span below and above human adult dose of silymarin. *P<0*.*05* *= Statistically significant

#### Survival Assay

The fly population was split into 5 groups of 50 young flies each (both sexes), with 5 replicates and a control group. For 28 days, the flies were exposed to fly food containing Silymarin powder in varying concentrations and flipped every 3 days to newly prepared diet for consistency. Until the experiment’s conclusion, the number of alive and dead flies were scored daily. The life/dead fly percentage score were expressed^10^(fig.2).

#### Locomotor Performance

Locomotor performance was conducted to investigate the effects on neuromuscular and skeletal muscle as described by Le Bourg and Lints^11^. After the treatment period, the locomotor performance of the control and treated flies were assessed using the negative geotaxis assay. Ten flies from each group—control and treated—were individually immobilized in cold ice anaesthesia using labelled vertical glass columns (15 cm long, 2 cm wide). The flies were tapped at the base of the column following 10 minutes of recovery, and the number of flies that passed the 6 cm line in 6 s were noted. Normally, locomotory-normal flies travel quickly to the top, but locomotory-defective flies move slowly and may hang around towards the bottom. The climbing scores represent the average proportion of the total number of flies in an experiment that reached the 6 cm line. The data was presented as the percentage of flies that escaped in three separate studies beyond a minimum distance of 6 cm in 6 seconds (fig. 5.).

**Fig 5:**
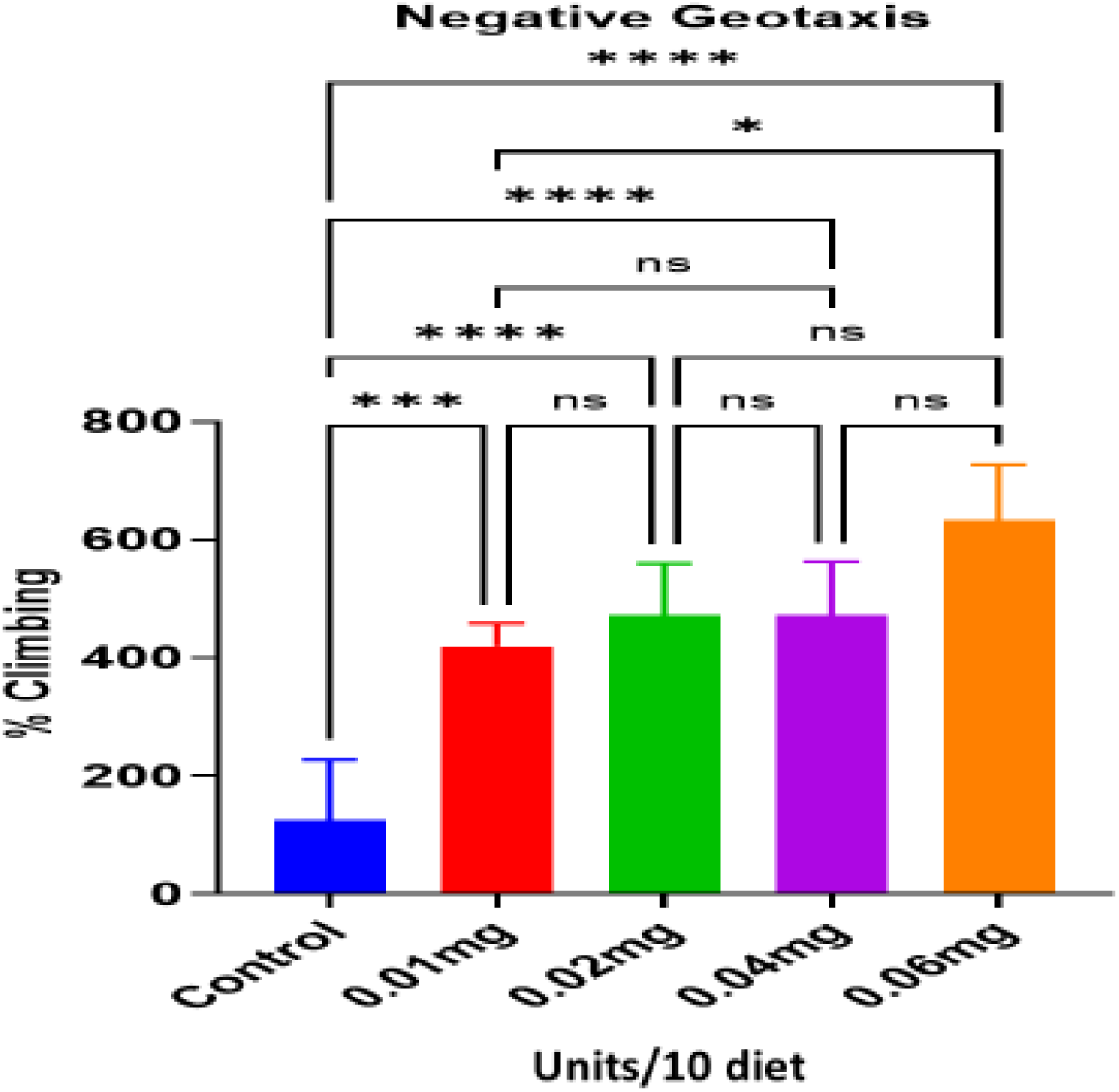
Showing Negative Geotaxis. This result accounts for a concentration-dependent increase (*P<0*.*05*) in the DM locomotor activity. The climbing rate has been used in DM as a measure of exercise and it describes the physical difficulty of the route. The increased climbing rate, therefore, signifies increased exercise and increased ability to overcome geotactic huddles. *P<0*.*05* *= Statistically significant

#### Estimation of Total Thiol Level

This is to assess the level of free radicals, signal transduction, apoptosis and various functions at the molecular level that may be due to the effects of the drug^12^. Total thiol was determined according to standard procedures as described by Elman^13^, but with some modifications. In brief, the reaction mixture included 20 µL of the sample, 170 µL of 0.1 M potassium phosphate buffer (pH 7.4), and 10 µL of 5,5′-dithiobis-(2-nitrobenzoic acid (DTNB). Using a microplate reader, the absorbance at 412 nm was measured following incubation for 30 min at room temperature. The material was precipitated with 4% sulfosalicylic acid in a 1:1 ratio for non-protein thiol. The samples were stored at 4 °C for 1 hour before being centrifuged for 10 minutes at 5000 rpm. The assay combination contained 10µL of DTNB, 20 µL of supernatant, and 170 µL l of 0.1 M phosphate buffer. The reaction was allowed to sit at room temperature for 30 minutes, and then a microplate reader was used to measure the reaction’s absorbance at 412 nm. Reduced Glutathione (GSH) was used as the reference for both total and non-protein thiols, and the data were represented as mol/mg of protein.

#### Determination of Acetylcholinesterase (AchE) Activity

AchE activity was determined according to standard methods Iorjiim, et al. ^10^ to investigate the level of acetylcholine and effects at the neuromuscular junction. Briefly, the Silymarin-treated flies were centrifuged using a cold centrifuge (Eppendorf AG, 5227 R, Germany) for four minutes at a speed of 4,000 rpm after being anaesthetized on ice and homogenized in 1:10 volumes of 100 mM phosphate buffer saline (pH 7.4). The method described by Elman ^13^ was followed, with few minor modifications, to collect the supernatant and use it to measure AChE activity. Sixty (60) µL of 8 mM acetylthiocholine was added to the reaction mixture, which also contained 285 µL of distilled water, 180 µL of 100 mM potassium phosphate buffer (pH 7.4), 60 µL of 10 mM DTNB, and 15 µL of sample. Using a UV spectrophotometer, the absorbance change was tracked at 412 nm for 2 min at 10 s intervals (Jenway Spectrophotometer 7315). The protein concentration of the whole fly homogenates was measured using the total protein kit (Randox) and following the manufacturer’s instructions. The results were adjusted by the protein content after the data were calculated against blank and sample blank. Micromole/min/mg of protein was used to express the enzyme activity.

#### Determination of Catalase (CAT) Activity

Catalase activity was measured spectrophotometrically using the Aebi ^14^ method by monitoring the disappearance of H2O2. The reaction medium was made up of 1800 µL of 50 mM phosphate buffer (pH 7.0), 180 µL of 300 mM H2O2, and 20 µL of the sample (1:10 dilution). A UV-visible spectrophotometer was used to measure the reaction for 2 minutes (10 s intervals) at 240 nm (Jenway Spectrophotometer 7315). CAT activity was measured in millimoles of H2O2 consumed per minute per milligram of protein.

#### Determination of Glutathione -S-Transferase (GST) Activity

Glutathione-S-transferase activity was measured using the Habig and Jakoby^15^ technique with 1-chloro-2,4-dinitrobenzene (CDNB) as the substrate. 270 µL of solution A (20 µL of 0.25 M potassium phosphate buffer, pH 7.0, with 2.5 mM EDTA, 10µL of distilled water, and 500 µL of 0.1 M GSH at 25 °C), 20 µL of the sample (1:5 dilution), and 10 µL of 25 mM CDNB comprised the test combination. In a SpectraMax microplate reader, the reaction mixture was monitored at 340 nm for 5 minutes at 10 s intervals The data were given in mol/mins/mg protein.

#### Fecundity Assay

Equal numbers (five each) of male and female Drosophila were exposed to graded concentrations of Silymarin for 5 days. Afterwards, the egg-laying pattern and eventual development were assayed for 14 days. The fertility of the flies following Silymarin administration was evaluated using the reproductive ability assay ^16^ with minor changes. In brief, virgin flies (both sexes) were isolated (within 8 hours of eclosion) from their regular fly diet and treated for five days with a series of Silymarin treatments (. All the flies in each treatment group were placed into fresh vials with a normal diet for 24 hours. The average number of emerging flies provides a measurement of morphological and reproductive capacity.

### Statistical Analysis

Data obtained were statistically analyzed using one-way ANOVA and multiple t-test of Graph pad Prism version 8 and the results were presented as mean ± standard deviation. A 95 % confidence level was used to determine the statistical difference between the control and the treated and between groups.

#### SARS-CoV-2 intro studies with Silymarin compound on Vero cell lines

##### In vitro experimental treatment with Silymarin

###### Cell culture and virus propagation

SARS-CoV-2 detected from oropharyngeal swabs by qRT-PCR were cultured in vero E6 monolayer cells (Vero E6, ATCC CRL-1586) following the protocol described by Leland and French (1988) and maintained in Dulbecco modified Eagle’s medium (DMEM) (Sigma, Germany) supplemented with 10% heat-inactivated fetal bovine serum (FBS) (Gibco, USA), 100 mM L-glutamine, 100 U/ml penicillin, 100 μg/ml streptomycin (Biological Industries, USA). Cells were maintained in a chamber incubator at 37°C under a 5% CO_2_ atmosphere saturated with >90% humidity. The virus was isolated, and titre was determined following Yao et al. (2020) and Reed and Muench method (Reed and Muench, 1938).

###### Viral infection and drug treatment

Subconfluent Vero cells were grown on ninety-six (96) well plates and infected with SARS-CoV-2. The study was divided into pre- and post-exposure for each compound concentration to be assessed. The pre-and post-treatment means the addition of either virus or compound first and a third group where the virus and compound were added to the Vero cells all at once. The virus or compound was allowed to adsorb for 30 min before adding the compound or virus respectively.

Different concentrations of Silymarin (**a**. S1-250ug/ml, **b**. S2-300ug/ml, **c**. S3-350ug/ml, **d**. S4-500ug/ml) were prepared and used to determine the pre or post-exposure activity of the compound against a fixed titre of the virus. Each concentration was run in quadruplicate. The experiment was observed for 5 days and activity was monitored under an inverted microscope for five days before being terminated and confirmed by RT-PCR. Viral RNAs were extracted from cultures using QIAamp Viral RNA Mini Kit (Qiagen, Germany) following the manufacturer’s instructions. The result was generated using GeneFinder™ COVID-19 Plus RealAmp Kit (Korea) targeting three genes namely: RdRp gene, N gene and E gene on a Thermo Scientific Piko Real Time-PCR system using the recommended cycling conditions (50 °C for 20 min, 95 °C for 5 min, followed by 45 cycles of 95 °C for 15 s and 58 °C for 60 s). The human housekeeping gene RNAse P was targeted as the internal control for optimal nucleic acid extraction and the absence of PCR inhibitors in all assessed samples. Cycle threshold (Ct) values of ≤ 40 and lower were considered positive, and values between ≥ 40 were considered negative.

## Results

The LD_50_ experimental results(Figure 1) showed that Silymarin is practically safe in the *Drosophila melanogaster* model. Statistical analysis showed LD_50_ range of above 12,000 mg/10g diet. There was no significant difference (P > 0.05) between the various groups and the control. The seven days acute toxicity tests demonstrate over 90% fly survival in all the groups, and there was no significant difference (*P>0*.*05*) between the treatments and the control (Figure 2). Twenty-eight (28) days survival assay of daily exposure to silymarin dose range between 50% to 2000% adult dose showed no adverse effects (*P>0*.*05*) on the fly population (Figure 3). These results show that chronic exposure to silymarin is equally safe with well above 75% survival 28 days post-exposure in all the groups. The dose of 0.01mg, 0.02mg, 0.04mg, and 0.06mg per 10g diet corresponding to 50%, 100%, 200%, and 500% AD was determined for seven days experiment (Figure 4).

### Locomotor Performance

There was a dose depended on increase (*P<0*.*05*) in the *Drosophila* ability to climb against gravity. A negative geotaxis experiment measures the *Drosophila* exercise level and ability to escape hurdles. The dose of 0.06mg/10g diet showed an increased (P=0.0001) in the fly climbing ability (Figure 5).

### Biochemical Parameters

*Drosophila melanogaster* exhibits increased levels of total protein and antioxidant thiols when exposed oxidative stress and toxicants. The results of the 7 days exposure to silymarin showed no significant difference in both the treated and the control groups (Figure 6&7). The results of the acetylcholinesrase (AChE) level showed no significant (*P>0*.*05*) change in the treated group compared with the control (Figure 8). Changes in acetylcholinesterase level is observed during neuromuscular junction toxicology in DM. Catalase is an early antioxidant enzyme mobilized by DM during exposure to toxicants. The result of the seven days continuous exposure to silymarin revealed no significant (*P>0*.*05*) difference between the silymarin and the control group (Figure 9). Glutathione-S-transferase is an important antioxidant enzyme that detoxify both endogenous and exogenous chemicals. Increased levels of GST is seen in DM in toxicity. The results of this experiment showed no change (*P>0*.*05*) in GST levels of all the groups (Figure10).

**Fig 6:**
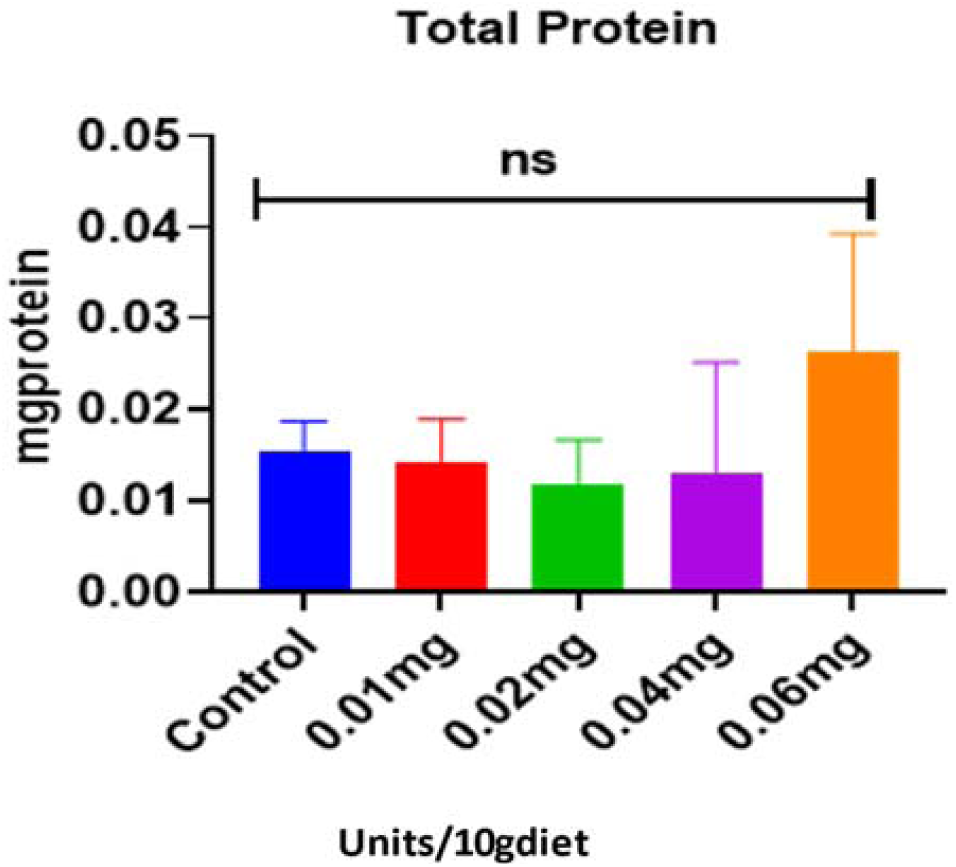
The Results Showed Total Protein Determination of 7-Days Oral Daily Dose of Silymarin in DM. There was no significant difference (*P>0*.*05*) between the total protein in the different groups and the control. Toxicants are known to result in increased total protein levels in the fly homogenate. *P<0*.*05* *= Statistically significant

**Fig 7:**
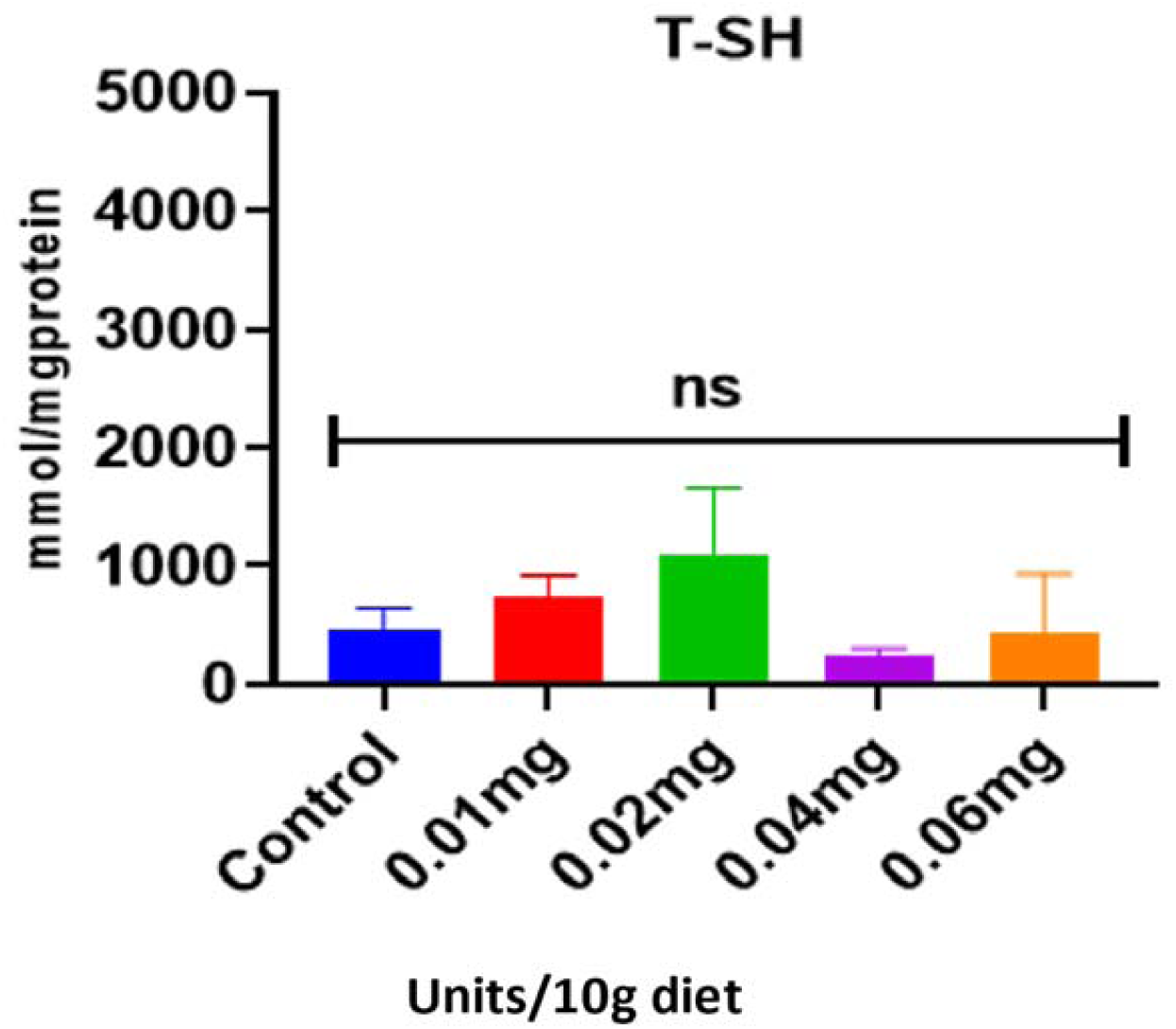
Showing the Total Thiol Levels of the DM after 7-Days oral Daily Dose. There was no statistical difference (P>0.05) between the treated and the control groups. *P<0*.*05* *= Statistically significant

**Fig 8:**
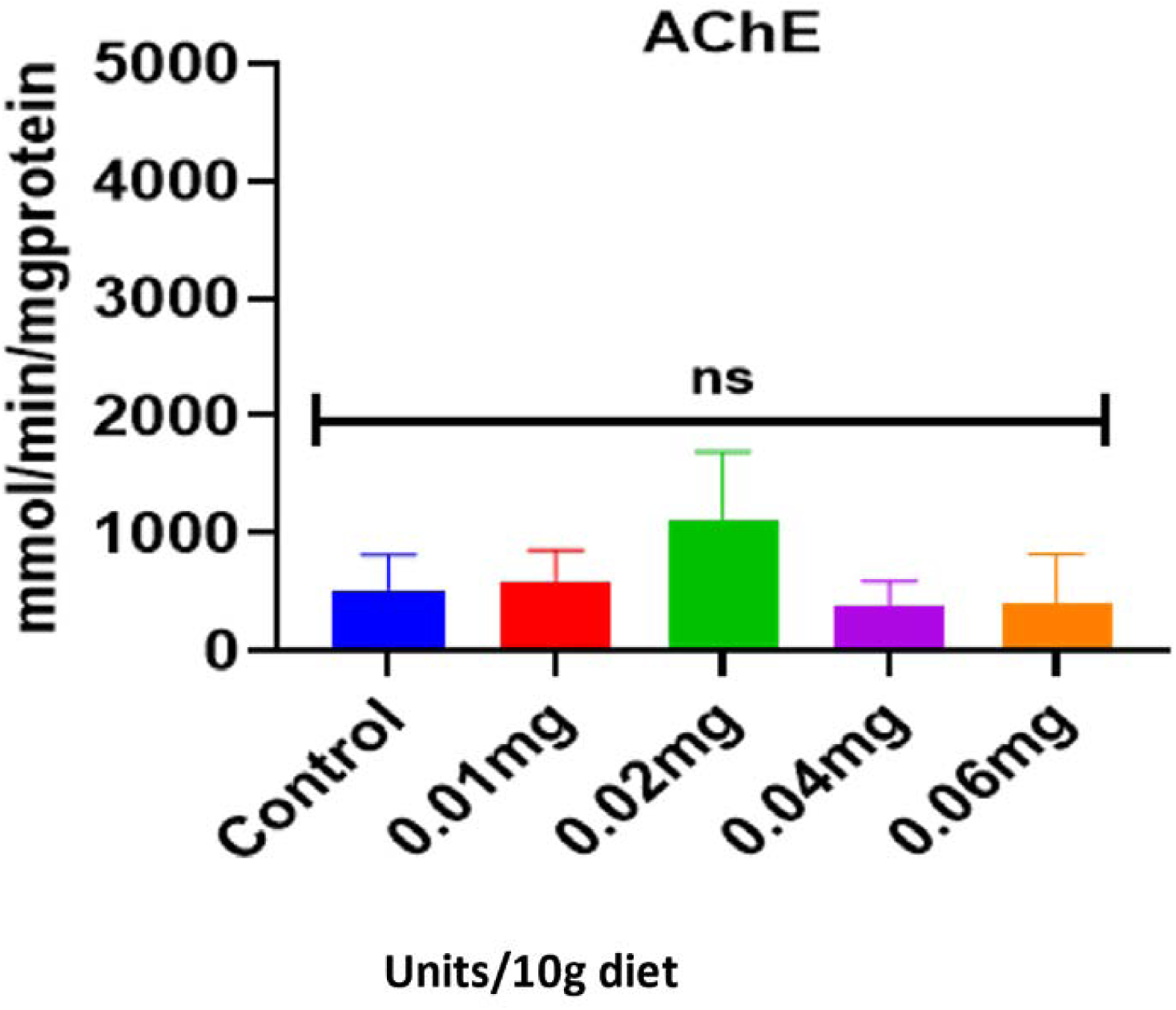
Showing the Fig: 8Acetylcholinesterase Levels of the DM after 7-Days oral Daily Dose. There was no statistical difference (P>0.05) between the treated and the control groups. *P<0*.*05* *= Statistically significant

**Fig 9:**
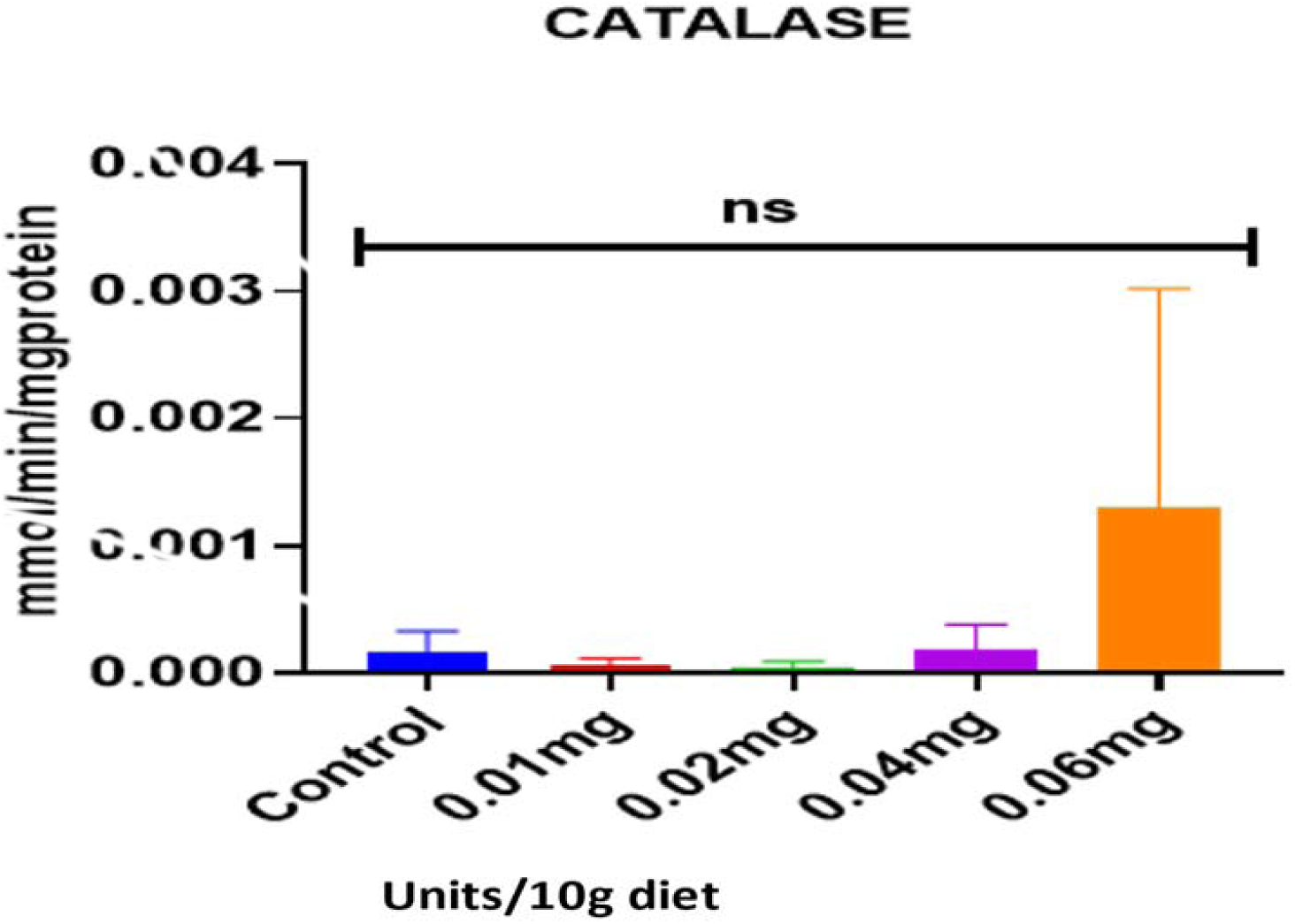
Showing Catalase Levels of the DM after 7-Days oral Daily Dose. There was no statistical difference (P>0.05) between the treated and the control groups. *P<0*.*05* *= Statistically significant

**Fig 10:**
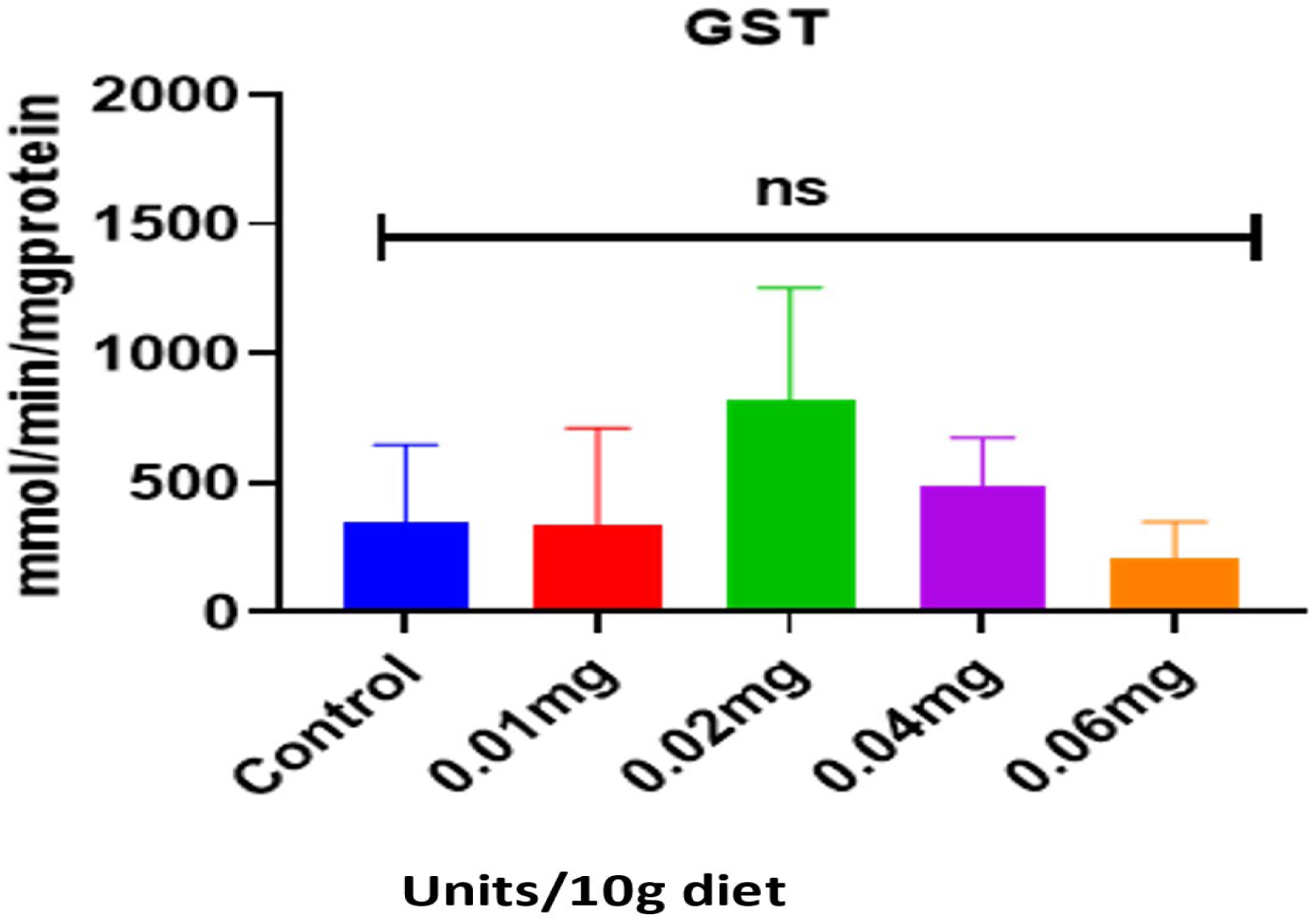
Showing the Glutathione-S-Transferase (GSH) Levels of the DM after 7-Days oral Daily Dose. There was no statistical difference (P>0.05) between the treated and the control groups. *P<0*.*05* *= Statistically significant

### Fecundity

The results of the *Drosophila* exposure to the graded concentration of silymarin exhibited no significant difference (*P>0*.*05*) in all the groups (Figure 11).

**Fig 11:**
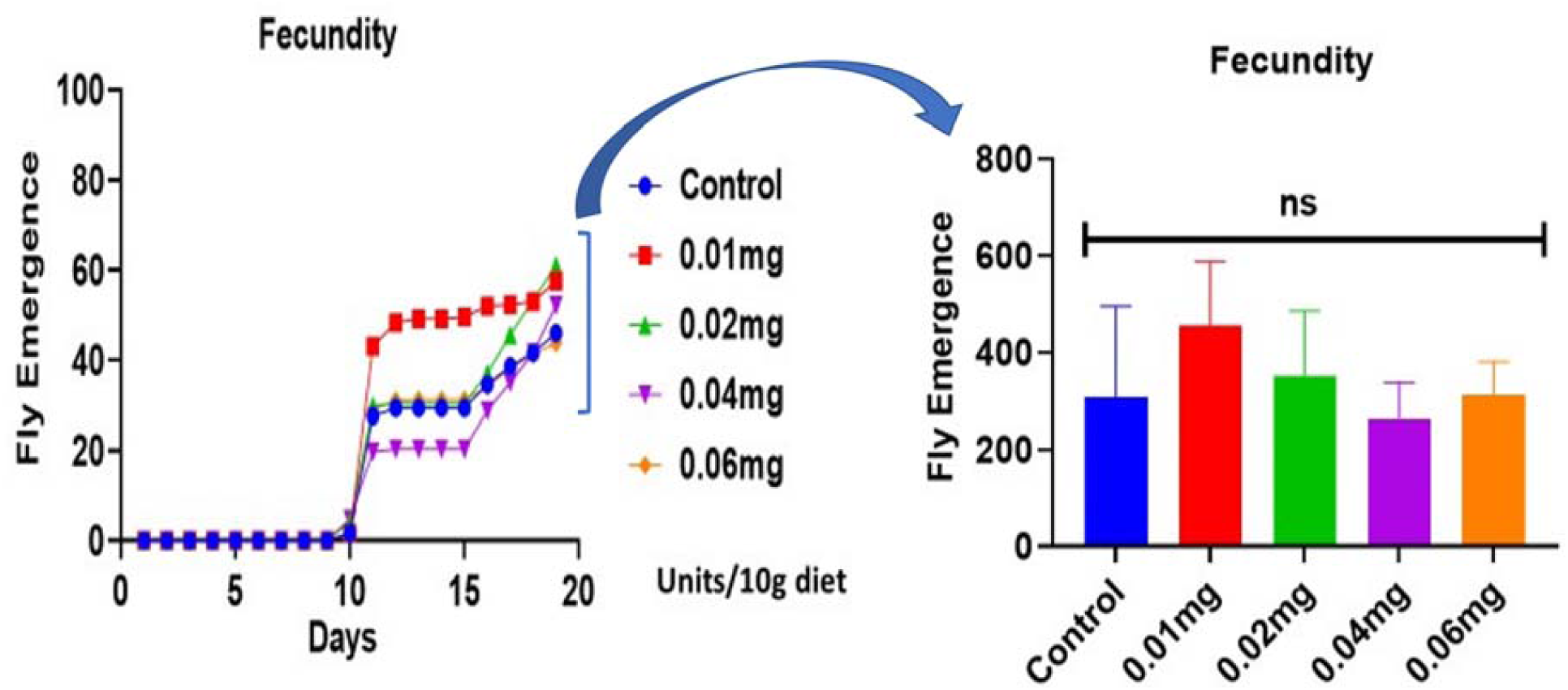
Showing the Adult Fly Emergence After 7-Days Oral Daily Dose of Silymarin. There was no statistical difference (P>0.05) between the treated and the control groups. However, more flies emerged cumulatively from the eggs of the treated groups (61%,57.6%, 52.5%) compared with the control (46%) *P<0*.*05* *= Statistically significant

**Figure 12:**
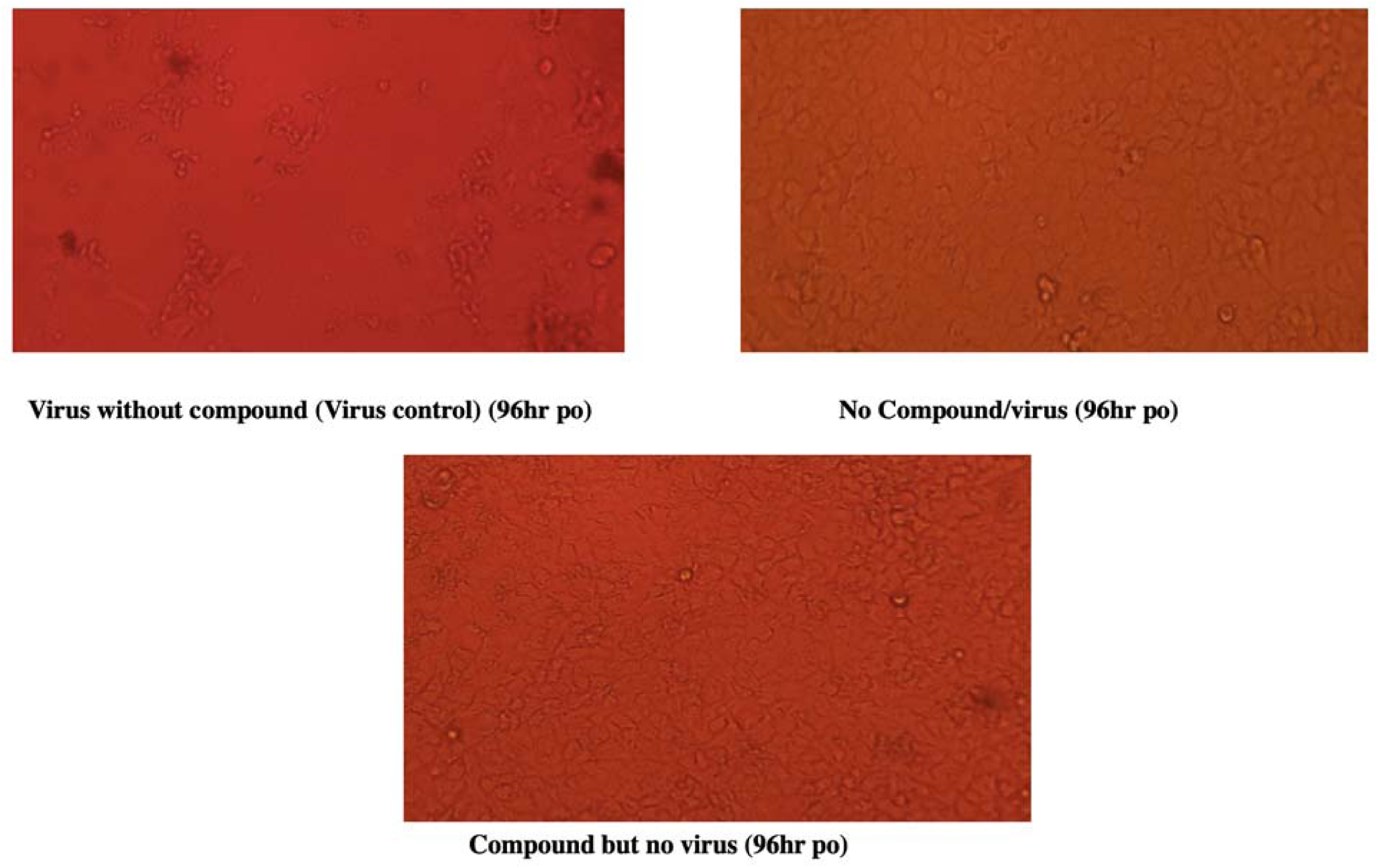
Different controls used for the study after 96 hours

**Figure 13:**
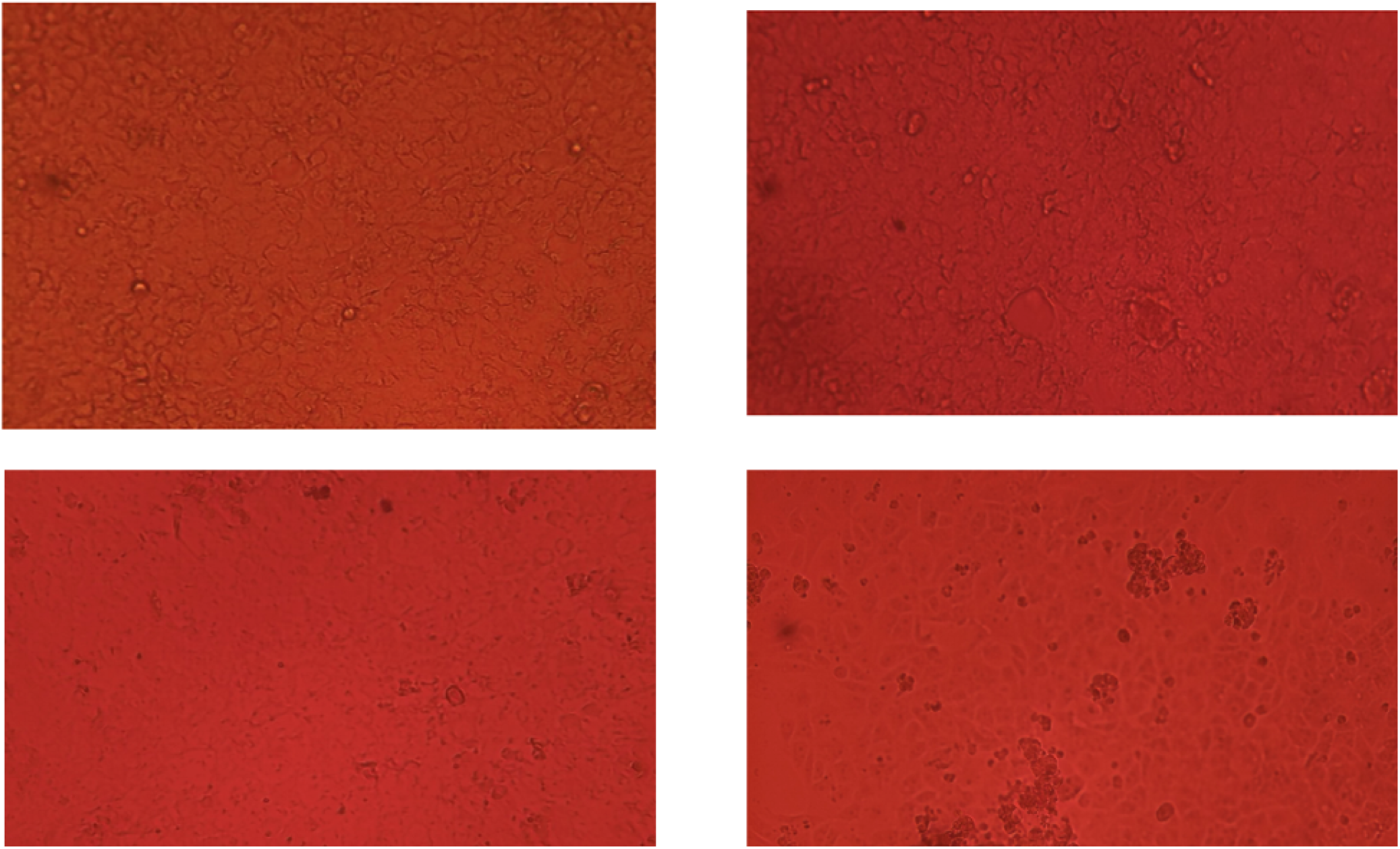
Pre-exposure S1pr_S4pr (250ug, 300ug, 350ug, 500ug) clockwise (48-96hr po)

**Figure 14:**
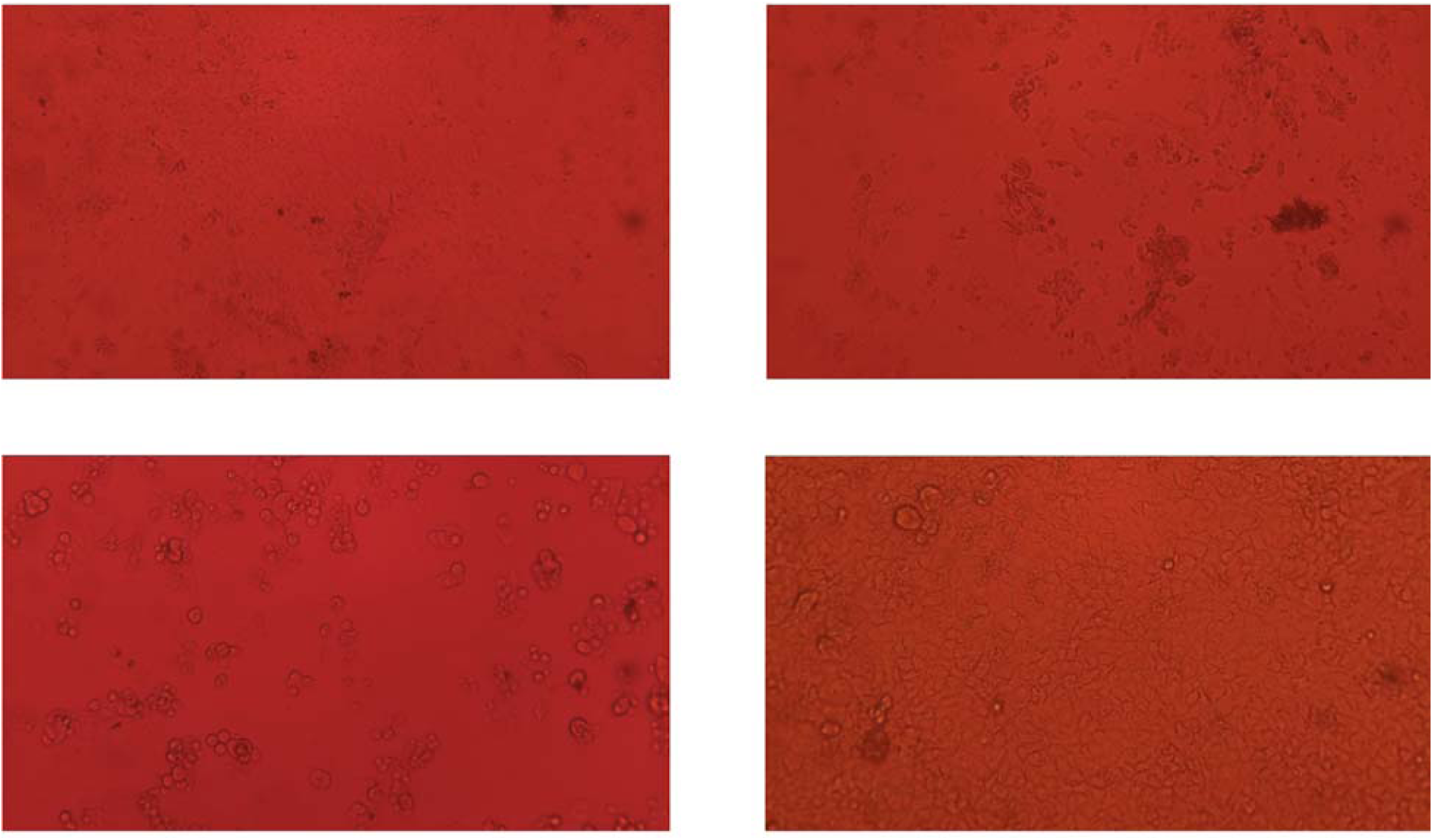
Post-exposure S1po – S4po (250ug, 300ug, 350ug, 500ug) clockwise (48-96hr po)

**Figure 15:**
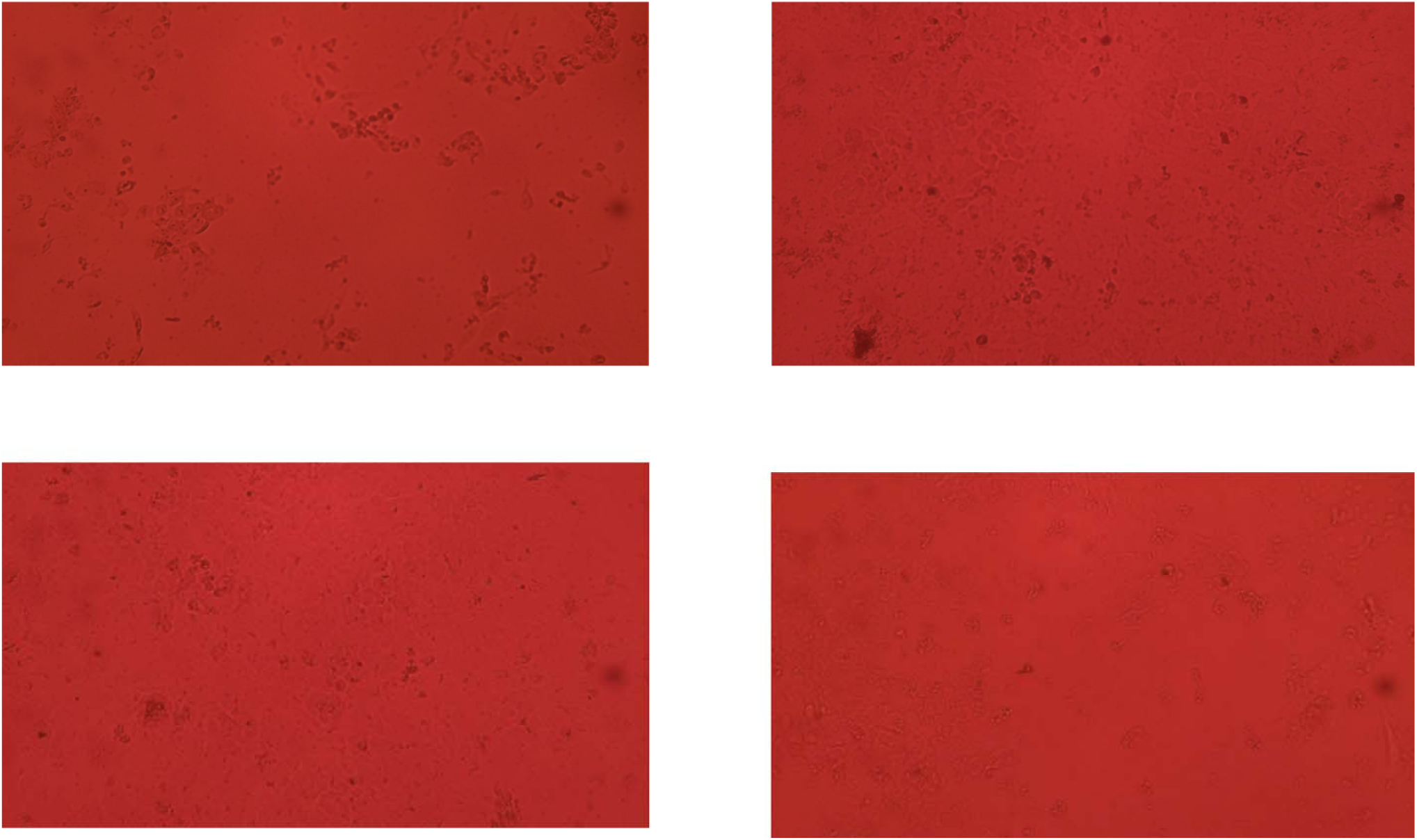
Concurrent exposure (S1s-S4s) (250ug, 300ug, 350ug, 500ug) clockwise (48-96hr po)

### Invitro experiment

The titre of the virus used for the experiment was determined as 10^4.5^ TCID50)/mL. However, an initial concentration of the compound (a. S1a-100 µg/ml b. S2a-45 µg/ml, c. S3a-11.25 µg/ml, d. S4a-0.703 µg/ml, e. S5a-0.176 µg/ml) was used for the experiment and within 48 hrs, rounding and detachment of cells from the surface of the plates were observed for both pre and post-exposure and for wells where the virus and compound were added at the same time across all the plates. A second dilution of the compound was prepared with higher concentrations: a. S1-250ug/ml, b. S2-300ug/ml, c. S3-350ug/ml, d. S4-500ug/ml) and its activity on the virus assessed.

For treatment group A (pre-treatment): wells S1pr-S4pr, (Compound Silymarin before SARS-CoV-2), all inoculated and incubated Vero cells in quadruplicate wells were observed under the inverted light microscope 24 and 48 hrs post-infection and showed no cytopathic effect (CPE) (Fig 1 D) like the control (Fig 1A, B) wells inoculated with plain media after 48 - 72 hrs. Similarly, treatments group B, S1po-S4po (SARS-CoV-2 before the compound Silymarin), All inoculated 24 and 48 hrs, revealed rounding and detachment of cells (Fig 1 C). In contrast, the control wells inoculated with plain media in duplicate were observed without CPE.

In the third group, treatment C, S1s-S4s12, (Compound Silymarin plus SARS-CoV-2 simultaneously), cytopathic effects characterized by rounding and detachment of cells were observed. The experimental treatment of Vero cells with Silymarin at the concentration of 250 - 500 ug/ml all revealed a pre-treatment effect to SARS-CoV-2 in vitro. All experiments with the virus first showed no inhibition of CPE to the different concentrations of the compound. Silymarin does not seem to affect cells treated with compound and virus simultaneously. All products of the three experimental groups were subjected to qPCR, and the Cq values ranged between 21.01 – 32.23 (post-treatment), 29.33 – 40.56 (pre-treatment), 20.25 – 21.43(virus/silymarin). The positive control gave a Cq value of 20.5 −22.73.

## Discussion

*Drosophila melanogaster* (fruit-fly) shared close physiological, genetic resemblance to humans with 60% homology, and about 75% of human disease genes are conserved in the fruit-fly^17^. *Drosophila* has been used to study the toxicity of xenobiotics ^18^. Daily exposure of DM to the graded concentration up to 2000% human adult dose of silymarin had no adverse effects during the treatment period (Figure 1), suggesting that silymarin has wide safety and therapeutic index. High LD_50_ (12,000mg/10g diet) indicates safety, while low LD_50_ indicates toxicity. Acute and chronic toxicity data are used to determine the safety of substances fed to *Drosophila*. Here we observed no statistical difference upon short or long-time exposure between the control and the treated groups. It could be inferred from these results that silymarin has no adverse effects on short-time or chronic therapy. Vahid et al. reported that silymarin is safe in humans at a dose of 700mg thrice daily for 24 weeks^19^

The climbing assay (Negative geotaxis) has been used to study the effects of chemicals on exercise, locomotion, learning, and memory in the *Drosophila melanogaster* ^*20,21*^. The increased climbing activity observed in this study indicates silymarin’s ability to improve learning and memory, locomotion, orientation, and location in DM after tapping to the bottom. Memory loss in diabetic rats was significantly improved with silymarin supplementation^22^. Silymarin has also been reported to reduce symptoms of Parkinson’s disease in mice ^23^.

The biochemical markers (Total protein, total thiols, Acetylcholinesterase, Catalase, Glutathione-S-transferase) of oxidative stress assessed exhibits no significant deviation from the normal. This result suggest further safety profile of silymarin ^24^ as supported by the LD_50_, acute, and chronic toxicity tests results. *Drosophila melanogaster* has been used for developmental assay^25^. We studied effects of silymarin on egg-laying, eclosion and adult emergence as a standard procedure for evaluating effects of xenobiotics in *Drosophila*^*26*^. There was no significant difference (*P>0*.*05*) in egg-laying, pupariation, eclosion and adult-fly emergence (Figure 11).

Silymarin showed antiviral activity against SARS-CoV-2 in agreement with the report of Ubani et al. ^3^. The activity observed was dependent on the mode of administration of the compound. Antiviral activity was observed for all the concentrations using CPE evaluation and qPCR for all three methods of administration. Our results demonstrated that silymarin reduced CPE in SARS-CoV-2-infected Vero E6 cells and qPCR showed higher Cq compared to post-treatment and co-treatment with silymarin and virus at the same time suggesting a delay in the SARS-CoV-2 replication.

Our results suggest that silymarin has more preventive activity against SARS-CoV-2 than curative in vitro. However, all the concentrations of the compound exhibited antiviral activity against the virus. The anti-coronavirus properties of silymarin should be further investigated in vivo. Finally, we suggest that the minimal cytotoxic concentrations of silymarin be assessed against potential antiviral activity to obtain precise results and ensure that the methodology does not affect the quality of the data.

However, the mode of action is not known. However, we postulate that silymarin interacts with S glycoprotein to prevent SARS-COV-2 from entering the cell. Hence will be a good preventive drug. Silymarin’s antiviral effects have been studied in viruses such as HCV, influenza A virus, mayaro virus (MAYV), chikungunya virus (CHIKV), and human immunodeficiency virus (HIV). Silymarin has antiviral activity against all of the viruses mentioned, preventing virus entry into host cells, genomic content replication, and viral protein gene expression ^27^. A review of the preventive capability of Silymarin is reviewed elsewhere^28^

## Conclusion

The results of these experiments revealed that silymarin exerts no observable adverse effects on the *Drosophila melanogaster* survival, oxidative enzymes, egg-laying and development at the dose range of 50%-2000% human adult dose tested. Silymarin has a more preventive than the curative effect on SARS-COV-2. However, the mechanism of prevention should be evaluated.

## Notes

### Competing Interest Statement

The authors have declared no competing interest.

